# Oligonucleotide Capture Sequencing of the SARS-CoV-2 Genome and Subgenomic Fragments from COVID-19 Individuals

**DOI:** 10.1101/2020.12.11.421057

**Authors:** Harsha Doddapaneni, Sara Javornik Cregeen, Richard Sucgang, Qingchang Meng, Xiang Qin, Vasanthi Avadhanula, Hsu Chao, Vipin Menon, Erin Nicholson, David Henke, Felipe-Andres Piedra, Anubama Rajan, Zeineen Momin, Kavya Kottapalli, Kristi L. Hoffman, Fritz J. Sedlazeck, Ginger Metcalf, Pedro A. Piedra, Donna M. Muzny, Joseph F. Petrosino, Richard A. Gibbs

**Affiliations:** *H*uman Genome Sequencing Center, Baylor College of Medicine, Houston, Texas, United States of America; Alkek Center for Metagenomics and Microbiome Research, Department of Molecular Virology and Microbiology, Baylor College of Medicine, Houston, Texas, United States of America; Department of Molecular Virology and Microbiology, Baylor College of Medicine, Houston, Texas, United States of America, USA; Pediatrics, Baylor College of Medicine, Houston, Texas, United States of America, USA

## Abstract

The newly emerged and rapidly spreading SARS-CoV-2 causes coronavirus disease 2019 (COVID-19). To facilitate a deeper understanding of the viral biology we developed a capture sequencing methodology to generate SARS-CoV-2 genomic and transcriptome sequences from infected patients. We utilized an oligonucleotide probe-set representing the full-length genome to obtain both genomic and transcriptome (subgenomic open reading frames [ORFs]) sequences from 45 SARS-CoV-2 clinical samples with varying viral titers. For samples with higher viral loads (cycle threshold value under 33, based on the CDC qPCR assay) complete genomes were generated. Analysis of junction reads revealed regions of differential transcriptional activity and provided evidence of expression of ORF10. Heterogeneous allelic frequencies along the 20kb ORF1ab gene suggested the presence of a defective interfering viral RNA species subpopulation in one sample. The associated workflow is straightforward, and hybridization-based capture offers an effective and scalable approach for sequencing SARS-CoV-2 from patient samples.

## Introduction

The COVID-19 pandemic has spread worldwide with alarming speed and has led to the worst healthcare crisis in a century. The agent of COVID-19, the novel SARS-CoV-2 coronavirus (family *Coronaviridae*), has a ~30 Kb positive-sense single-stranded RNA genome predicted to encode ten open reading frames (ORFs) [1]. Similar to other RNA viruses, coronaviruses undergo mutation and recombination [2, 3] that may be critical to understanding physiological responses and disease sequelae, prompting the need for comprehensive characterization of multiple and varied viral isolates.

To date, reports highlighting genomic variation of SARS-CoV-2 have primarily used amplicon-based sequencing approaches (e.g., ARTIC) [4–7]. Attaining uniform target coverage is difficult for amplicon-based methods, and is exacerbated by issues of poor sample quality [8]. Genome variation in the amplicon primer region may also impact sequence assembly. Transcriptome characterization can further contribute to our knowledge of mutation within the SARS-CoV-2 genome, and direct RNA long read sequencing, both alone and in combination with short read sequencing, have been described [1, 9, 10]. Unfortunately, these analyses are equally hampered by sample quality limitations and necessitate use of cultured cell lines. Oligonucleotide capture (‘capture’) mitigates these issues as hybridization to specific probes not only enriches for target sequences but enables the analysis of degraded source material [11–14]. Capture enrichment has also been applied to viral sequencing, where a panvirome probe design resulted in up to 10,000-fold enrichment of the target sequence and flanking regions [15–17]. Direct RNA enrichment method has also been reported for viral genome sequencing, but each sample was enriched separately followed by pooling for sequencing [18]. Hybridization-based enrichment of RNAs can also aid in the identification of gene fusions or splice variants [13, 19, 20], which are particularly important for coronavirus biology. In addition to encoding a polyprotein that undergoes autocatalyzed hydrolysis, coronaviruses employ subgenomic RNA fragments generated by discontinuous transcription to translate proteins required for viral replication and encapsidation. These subgenomic RNA fragments share a common 62-bp leader sequence derived from the 5’ end of the viral genome, detectable as a fused junction to interior ORFs [1, 10]. Direct RNA sequencing of cultured cell lines infected with SARS-CoV-2 revealed that the junctional sequences are not evenly distributed between the ORFs, suggesting that individual proteins may be translated at different rates [1]. How virus translation profiles from infected human patients differ from those from cultured cells is as yet unknown.

Here we have utilized capture probes and a streamlined workflow for sequence analysis of both the SARS-CoV-2 genomic sequences and of the junction reads contained within the genomic subfragments generated by discontinuous transcription (Fig1). The method can be applied at scale to analyze samples from clinical isolates. Enriching for genomic and transcriptional RNA, followed by deep short-read sequencing, sheds light on variation in clinical SARS-CoV-2 genomic sequences and expression profiles.

**Fig 1.**
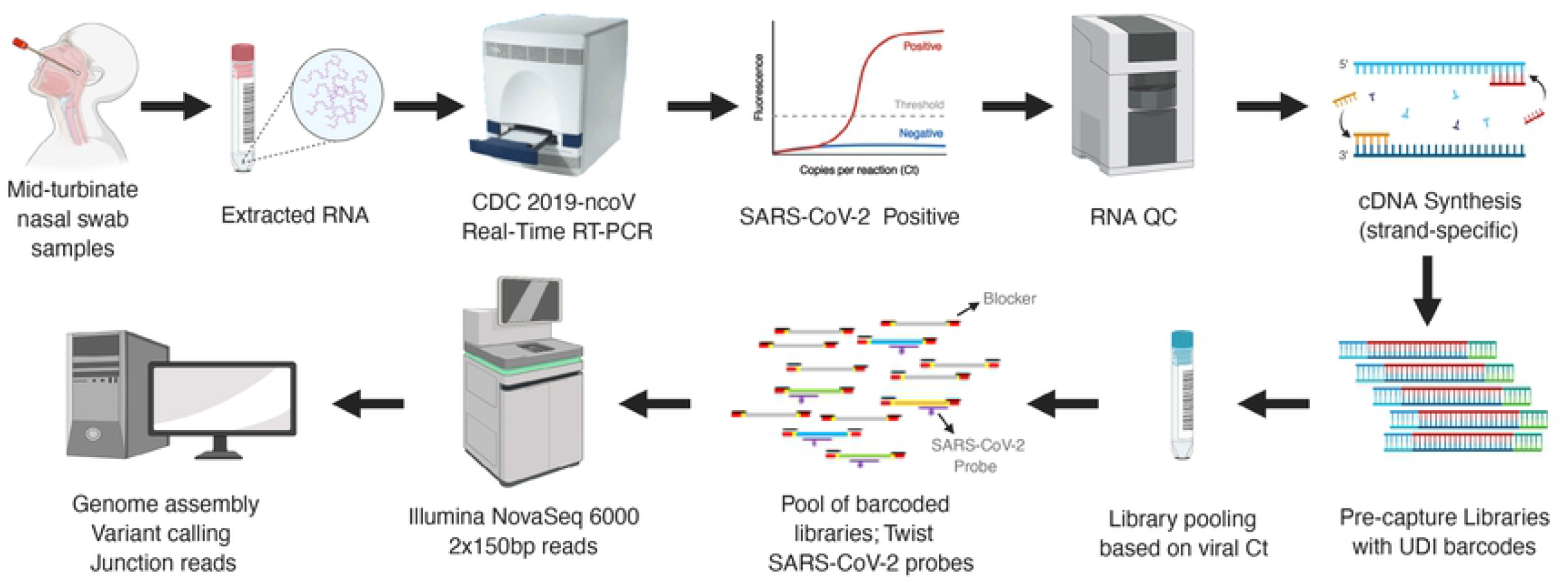
Schematic workflow. Presented in the workflow are the different steps involved in the SARS-CoV-2 capture and sequencing methodology.

## Material and methods

### COVID-19 viral testing, Collection, RNA extraction and real-time reverse transcription polymerase chain reaction (RT-PCR)

The CLIA Certified Respiratory Virus Diagnostic Laboratory (ID#: 45D0919666) at Baylor College of Medicine performed real time reverse transcriptase polymerase chain reaction (RT-PCR) tests for SARS-CoV-2 on mid-turbinate nasal swab samples collected from adults presenting to the hospitals or clinics at the Texas Medical Center from March 18 through April 25, 2020. Viral RNA was extracted from nasal swab samples using the Qiagen Viral RNA Mini Kit (QIAGEN Sciences, Maryland, USA) with an automated extraction platform QIAcube (QIAGEN, Hilden, Germany) according to the manufacturer instructions. Starting with 140 ul of the collected sample, nucleic acids were extracted and eluted to 100ul. All samples were tested by CDC 2019-Novel coronavirus (2019-ncoV) Real-Time RT-PCR Diagnostic panel. Primers and probes targeting the SARS-CoV-2 nucleocapsid genes, N1 and N2, were used. Samples were also tested for Ribonuclease P (RNase P) gene, to determine the quality of sample obtained. PCR reaction was set up using TaqPath™ 1-Step RT-qPCR Master Mix, CG (Applied Biosystems, CA) and run on 7500 Fast Dx Real-Time PCR Instrument with SDS 1.4 software. Samples with cycle threshold (Ct) values below 40 for both SARS-CoV-2 N1 and N2 primers were necessary to determine positivity. For seven samples with very low viral loads (N=7); Ct >37 and <40, the RNA was concentrated 4-fold by doubling the extraction volume - 280 μl and halving the elution volume - (50 μl) and submitted for sequencing.

### Library, capture, sequencing

#### Sequenced samples

Forty-five mid-turbinate nasal swab samples were collected from 32 unique individuals (S1 Table). The RNA Integrity Number (RIN) values ranged from 2.3 and 5.2 with Ct values from 16-39. The amount of RNA used as input for cDNA varied from 13.6 ng to 120 ng (S1 Table). As positive controls, 1,500 (Ct=36.2) and 150,000 copies (Ct=29.6) of the Synthetic SARS-CoV-2 RNA from Twist Biosciences (Cat# 102024) were spiked into two 50 ng Universal Human Reference RNA samples. To generate the synthetic RNA, six non-overlapping 5 Kb fragments of the SARS-CoV-2 reference genome (MN908947.3) sequence were synthesized by Twist Inc. as double stranded DNA, and transcribed in vitro into RNA. Three SARS-CoV-2 free mid-turbinate nasal swab samples which were negative for SARS-CoV-2 by real-time RT-PCR, were sequenced as negative controls. Due to limited sample size in this study, no other patient metadata was used to interpret results.

#### cDNA Preparation

cDNA was generated utilizing NEBNext^®^ RNA First Strand Synthesis Module (E7525L; New England Biolabs Inc.) and NEBNext^®^ Ultra™ II Directional RNA Second Strand Synthesis Module (E7550L; NEB). Total RNA in a 15 μl mixture containing random primers and 2X 1^st^ strand cDNA synthesis buffer were incubated at 94°C for 10 min to fragment the RNA. RNA were converted to cDNA by adding a 5-μl enzyme mix containing 500ng Actinomycin D (A7592, Thermo Fisher Scientific), 0.5μl RNase inhibitor, and 1 μl of Protoscript II reverse transcriptase, then incubated at 25°C for 10 minutes, 42°C for 50 minutes, 70°C 15 minutes, before being cooled to 4°C on a thermocycler. Second strand cDNA were synthesized by adding a 60 μl of mix containing 48 μl H_2_O, 8 μl of 10X reaction buffer, and 4 μl of 2^nd^ strand synthesis enzyme, and incubated at 16°C for 1 hour on a thermocycler. The double strand (ds) cDNA were purified with 1.8X volume of AMPure XP Beads (A63882, Beckman) and eluted into 42 μl 10 mM Tris buffer (Cat#A33566, Thermo Fisher Scientific). Because these libraries were prepared primarily for sequence capture, rRNA depletion or Ploy A+ isolation steps were not performed.

#### Library preparation

The double-stranded cDNA was blunt-ended using NEBNext^®^ End Repair Module (E6050L, NEB). Five μl 10X ER reaction buffer and 5 μl ER enzyme were added to the ds cDNA. The ER reaction was incubated for 30 minutes at 20°C on a thermocycler. After the ER reaction, cDNA were purified with 1.8X volume AMPure XP Beads and eluted into 42 μl nuclease free water (129114, Qiagen). Next, 5 μl of 10X AT buffer and 3 μl of Klenow enzyme from NEBNext^®^ dA-Tailing Module (E6053L, NEB) was added to the sample. The AT reaction was incubated at 37°C for 30 minutes. After incubation, samples were purified with 1.8X volume AMPure XP Beads and eluted into 33 μl nuclease free water (129114, Qiagen). Illumina unique dual barcodes adapters (Cat# 20022370) were ligated onto samples by adding 2 μl of 5uM adapter, 10 μl 5X ligation buffer and 5 μl of Expresslink Ligase (A13726101, Thermo Fisher), and incubated at 20°C for 15 minutes. After adapter ligation, libraries were purified twice with 1.4X AMPure XP and eluted into 20 μl H_2_O. Libraries were amplified in 50 μl reactions containing 150 pmol of P1.1 and P3 primer and Kapa HiFi HotStart Library Amplification kit (Cat# kk2612, Roche Sequencing and Life Science). The amplification was incubated at 95°C for 45 seconds, followed by 15 cycles of 95°C for 15 sec, 60°C 30 seconds, and 72°C 1 minutes, and 1 cycle at 72°C for 5 minutes. The amplified libraries were purified with 1.4X AMPure XP Beads and eluted into 50 μl H_2_O. The libraries were quality controlled on Fragment Analyzer [using DNA7500 kit (5067-1506, Agilent Technologies). The library yields were determined based on 200-800-bp range.

#### Capture enrichment and sequencing

cDNA libraries with Illumina adaptors constructed from SARS-CoV-2 positive individuals were pooled into six groups (S1 Table). Pools 1 and 2 were from batch 1 and pools 3-6 are from batch 2. The RT-qPCR Ct value of virus N gene varied in these pools as follows: Pool 1 with 6 samples (Ct 20.4 - 28.34); Pool 2 with 5 samples (Ct 29.75 - 37.95; Pool 3 with 5 samples (Ct 17.3 – 38); Pool 4 with 6 samples (Ct 27.8 - 39.3); Pool 5 with 11 samples (Ct 33 - 38.9) and Pool 6 with 12 samples (Ct 32.9 - 39.5). Pooled cDNA pre-capture libraries were hybridized with probes from the SARS-CoV-2 Panel (Twist, Inc) at 70°C for 16 hours. Total probe length is 120 Kb. Post-capture insert molecules were further amplified (12-16 cycles) to obtain the final libraries that were sequenced on Illumina NovaSeq S4 flow cell, to generate 2×150 bp paired-end reads. To evaluate the effect of hybridization-based enrichment 9 samples were sequenced before and after capture enrichment. Ribosomal RNA was removed computationally.

### Data analysis

#### Sequence Mapping, genome reconstruction and variant calling

Raw fastq sequences were processed using BBDuk (https://sourceforge.net/projects/bbmap/; BBMap version 38.82) to quality trim, remove Illumina adapters and filter PhiX reads. Trimming parameters were set to a k-mer length of 19 and a minimum Phred quality score of 25. Reads with a minimum average Phred quality score below 23 and length shorter than 50 bp after trimming were discarded. The trimmed fastqs were mapped to a combined PhiX (standard Illumina spike in) and human reference genome (GRCh38.p13; GCF_000001405.39) database using a two-step BBTools approach (BBMap version 38.82). Briefly, the trimmed reads were first processed through the bloomfilter script, with a strict k=31 to remove reads identified as human. The remaining reads were mapped to the reference genome with BBMap using a k-mer length of 15, the bloom filter enabled, and fast search settings in order to determine and remove hg38/PhiX reads. Trimmed and human-filtered reads were then processed through VirMAP [21] to obtain full length reconstruction of the SARS-CoV-2 genomes. SPAdes assembler [22] was also used for genome reconstruction. The resulting assemblies were compared to those from VirMAP. A reconstructed genome with >99% the length of the SARS-CoV-2 reference genome, NC_045512.2, was considered a fully reconstructed genome. Plots were generated using R (version 3.6.1) and the tidyverse (version 1.3.0) and ggplot2 (version 3.2.1) packages. Alignments and reference mapping were done using mafft [23] (version 1.4.0) and BBMap (version 38.82). Sequence variation compared to SARS-CoV-2 reference genome was performed using the genome alignment from mafft with in-house scripts. For heterozygous variant analysis, the sequence reads were aligned to the reference genome using BWA-mem [24] with default parameters, realigned using GATK [25], and variants were called using Atlas-SNP2 [26]. Variant annotation was performed with SnpEff [27] Lineage assignment of SARS-COV-2 following Rambaut et al (2020) used the Pangolin COVID-19 Lineage Assigner (https://pangolin.cog-uk.io).

#### Subgenomic mRNA and junction reads analysis

Illumina sequence reads were aligned to SARS-CoV-2 reference genome NC_045512.2 using STAR aligner v2.7.3a [28] with penalties for non-canonical splicing turned off as described by Kim et al^1^. Alignment bam files were parsed using an in-house script to obtain junction-spanning reads that contained the leader sequence (5’ end of the junction falls within 34 to 85 bp of the reference genome). Sub genomic RNAs were categorized by junction reads according to the genes of the immediate start codon downstream of the 3’ of the junctions. Junction read counts were normalized to the total number of mapped reads.

## Results

A total of 45 samples collected from 32 patients between March 18 and April 25, 2020 in Houston, Tx, USA were analyzed. These were a subset of individuals tested for the presence of SARS-CoV-2 early during the pandemic. RNA fractions were isolated from viral transport media and converted to cDNA. SARS-CoV-2 cDNA libraries were pooled into six groups (S1 Table). All 45 capture-enriched and nine of the pre-capture libraries were sequenced on an Illumina platform based on details provided in the online methods. A schematic workflow is shown in (Fig 1).

### Sequencing results and capture enrichment efficiency

A total of 7.15 billion raw reads were generated for the 45 SARS-CoV-2 positive samples sequenced (S1 Table). Since this study was to optimize the methodology, samples were sequenced deeper to ensure that results among samples were not biased. Sequences were trimmed to filter low quality reads and subsequently mapped to the GRCh38 reference genome to identify human reads (Fig 2A). Trimmed non-human sequence reads were analyzed using the VirMAP [21] pipeline where between 7-86.4% of total reads from post-capture libraries mapped to the SARS-CoV-2 reference. One sample (192000446B), which had only 6.37 ng total RNA starting material, did not generate any SARS-CoV-2 reads. Overall, the percentage of reads represented by SARS-CoV-2 was higher in samples with CDC protocol-based RT-qPCR Ct values <33 (Fig. 2A).

**Fig 2.**
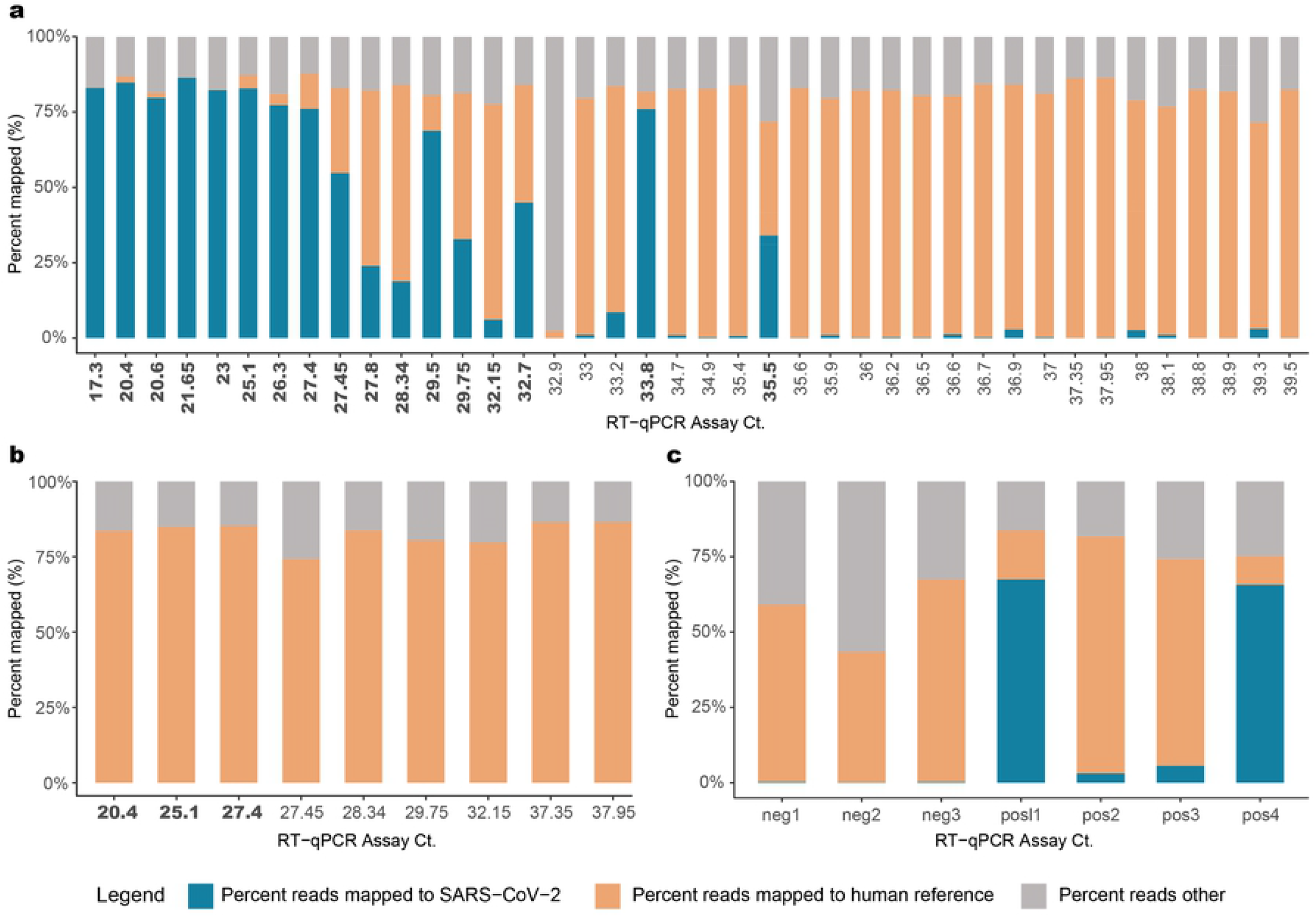
Sequence data. Ct value vs percent raw sequencing reads mapped to SARS-CoV-2 in (**a**) Capture enriched samples; (**b**) Pre-capture samples; (**c**) Positive and negative controls. Percentage of reads mapped to the ‘SARS-CoV-2’ genome, to the ‘human’ reference genome and a third category called the ‘reads others’, which is the combined total of trimmed reads and reads that do not fall under the two other categories are plotted in this figure. CT values in bold indicate samples that provided full-length genome assemblies.

To estimate the capture enrichment efficiency, pre-capture libraries for nine samples, ranging in Ct values of 20.4 to 37.95 (i.e. high to low titer in the original samples), were also sequenced, generating 152.1 – 322.9 million reads per sample. Samples 192000106B and 192000090B, with Ct > 37 produced zero reads mapping to the SARS-CoV-2 reference genome. In the remaining seven samples, less than 0.022% of reads were deemed SARS-CoV-2 (Fig 2B). Collectively, post-capture enrichment increased the SARS-CoV-2 mapping rate to 50.9%, a 9,243-fold enrichment.

Spiked synthetic SARS-CoV-2 RNA, encompassing six fragments of 5 Kb each, served as a positive control and were enriched successfully at both 1,500 and 150k copies per sample (Fig 2C). In the 1,500 copy libraries (n=2), 3-5% of reads mapped to the SARS-CoV-2 genome, while approximately 65% of reads from the 150k copy libraries (n=2) did the same (S1 Table). This translates to an approximate 91,858-fold enrichment in the 1,500 copy libraries and 13,778-fold enrichment in the 150k copy libraries compared to their starting amounts in the RNA. Three SARS-CoV-2 PCR negative samples were also sequenced, where <0.5% of reads mapped to the SARS-CoV-2 reference genome at 3-5 locations that are not conserved in the SARS-CoV-2 genome (S1 Table; S1 Fig).

### Genome reconstruction and genomic variations

In order to assess the ability of the capture methodology to assemble full-length genomes, both the nine pre-capture and 45 post capture libraries were assembled using both the VirMAP pipeline and the SPAdes *de novo* assembler_[22]^20^.

Full-length SARS-CoV-2 genomes were obtained from 17 of the 45 capture-enriched samples. Genome coverage in these 17 samples varied from 1071x to 3.19x million (S1 Table). Successful full-length genome assembly was correlated with Ct values below 33 (Fig 3), regardless of the total reads generated during sequencing. No variability between samples due to random priming of the cDNA synthesis or no gaps in genome coverage were noticed using this method (S2 Fig) Two samples with Ct values above 33, 192000296 (Ct 33.9) and 192000354 (Ct 35.5), obtained from a single patient, also yielded full-length genome reconstructions with acceptable quality (N ≤ 0.5%). Partial genome reconstructions were achieved for the remaining samples although somewhat surprisingly, the correlation between percentage of the genome that was reconstructed and the Ct value of that sample was not tightly correlated when Ct values were above 33 (Fig 3). Full-length genome sizes of the 17 capture-enriched and assembled sequences varied from 29.68 Kb to 30.15 Kb (S3 Fig). Variants relative to the SARS-CoV-2 reference genome sequence NC_045512.2, including single nucleotide polymorphisms and a single indel, ranged from 5 to 15 per sample, with a mean of nine.

**Fig 3.**
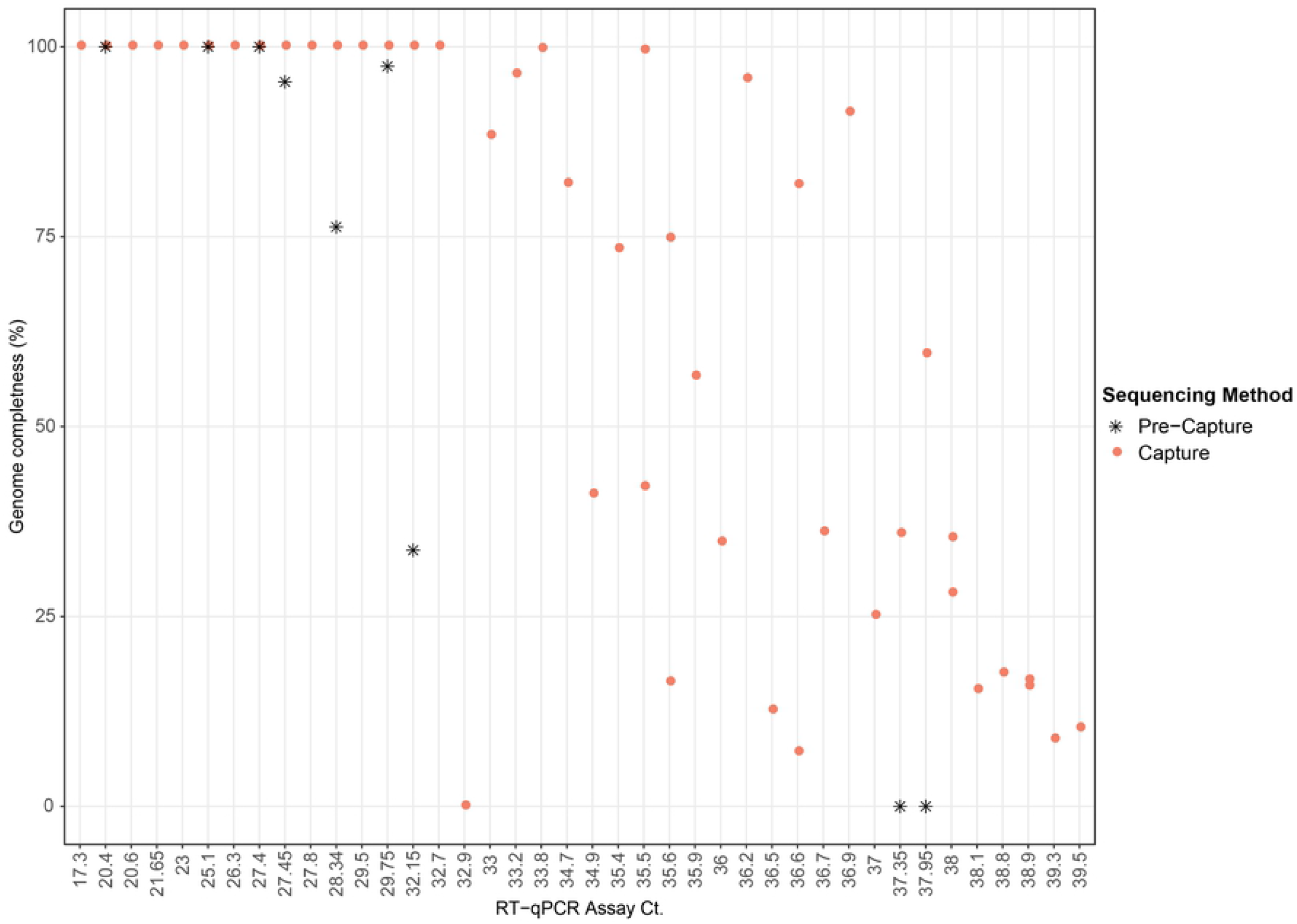
Scatter plot showing genome completeness as a function of Ct value. Pink circles represent post-capture samples and black asterisks represent pre-capture samples.

Out of the nine pre-capture samples, three (192000072B, 192000021B, 1920000003B), all with Ct values ≤ 27.4, yielded full-length genomes with 28x – 265x genome coverage, while in the other four samples, genome reconstructions were partial and also had a poor genome coverage of 1-6x. SARS-CoV-2 reads were not detected in the two remaining samples.

Alignment of DNA sequence reads from one sample (192000051B) to the reference SARS-CoV-2 genome sequence NC_045512.2 that is based on the first published isolate from Wuhan SARS-CoV-2 reference genome, revealed multiple heterogenous alleles (Fig 4; S4 Fig). Most isolates spreading into Europe derive from the ‘B’ lineage (based on the Wuhan sequence), but three samples including this sample contained an additional fraction of reads representing the A lineage [29] (S2 Table). Further investigation of the clinical correlates of this observation are underway. The genomic position 23,403 in the Wuhan reference strain had good coverage in 28 of the capture enriched samples. The A-to-G nucleotide mutation at this location that results in the Spike protein D614G amino acid change was noticed in 23 of the 28 samples [30].

**Fig 4.**
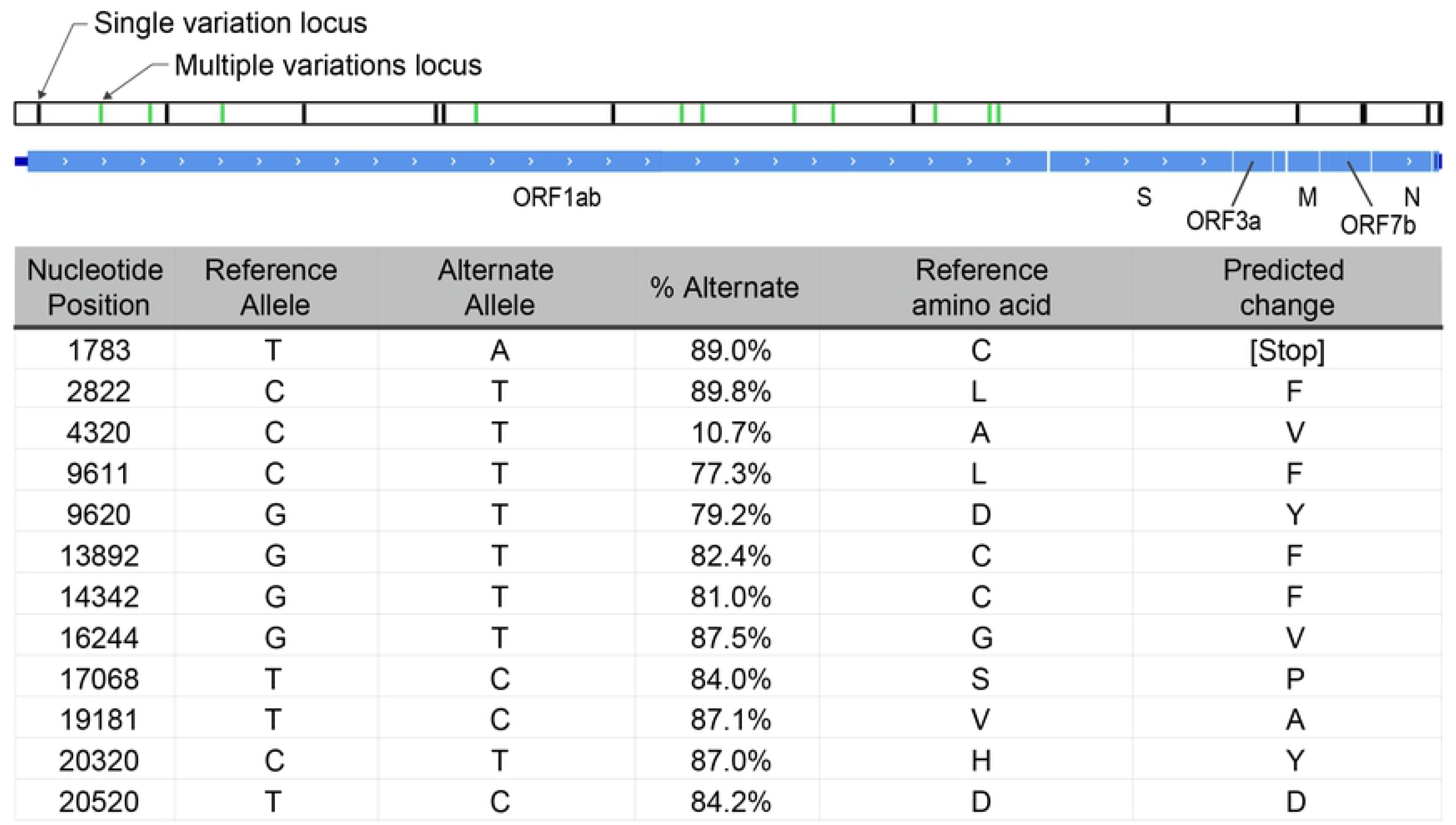
Schematic representation of 192000051B assembly. Black bars represent loci where the assembly called alleles different from the NCBI reference sequence NC_045512. Green bars represent mixed loci where both reference and alternative alleles were called. All mixed loci are in the ORF1ab gene, and are listed in the table, along with the frequency of the alternate allele at the position, and the predicted effect in translation.

### Characterization of SARS-CoV-2 subgenomic mRNAs

To identify and quantitate subgenome-length mRNAs, reads were aligned to the SARS-CoV-2 reference genome NC_045512.2. Only samples with full-length genomes (N=17 capture and N=5 pre-capture) were analyzed for junction reads to avoid introduction of any bias in identifying subgenomic RNA due to gaps in sequence coverage (Fig 5A and S3 Table). While full-length genomes were reconstructed from three pre-capture samples, an additional two samples with >95% genomes reconstructed, 192000135B (with 97.4%) and 192000088B (95.3%), were also included in this comparison (Fig 5A and in S4 Table).To characterize ORF expression in the capture and pre-capture libraries, the number of junction reads/million were calculated and plotted in Fig 5A (see details in S3 Table). Among the five pre- and post-capture comparison pairs, junction reads were identified in more ORFs after capture, and in instances where junction reads were found before and after capture, the expression trend agreed between the two groups.

**Fig 5.**
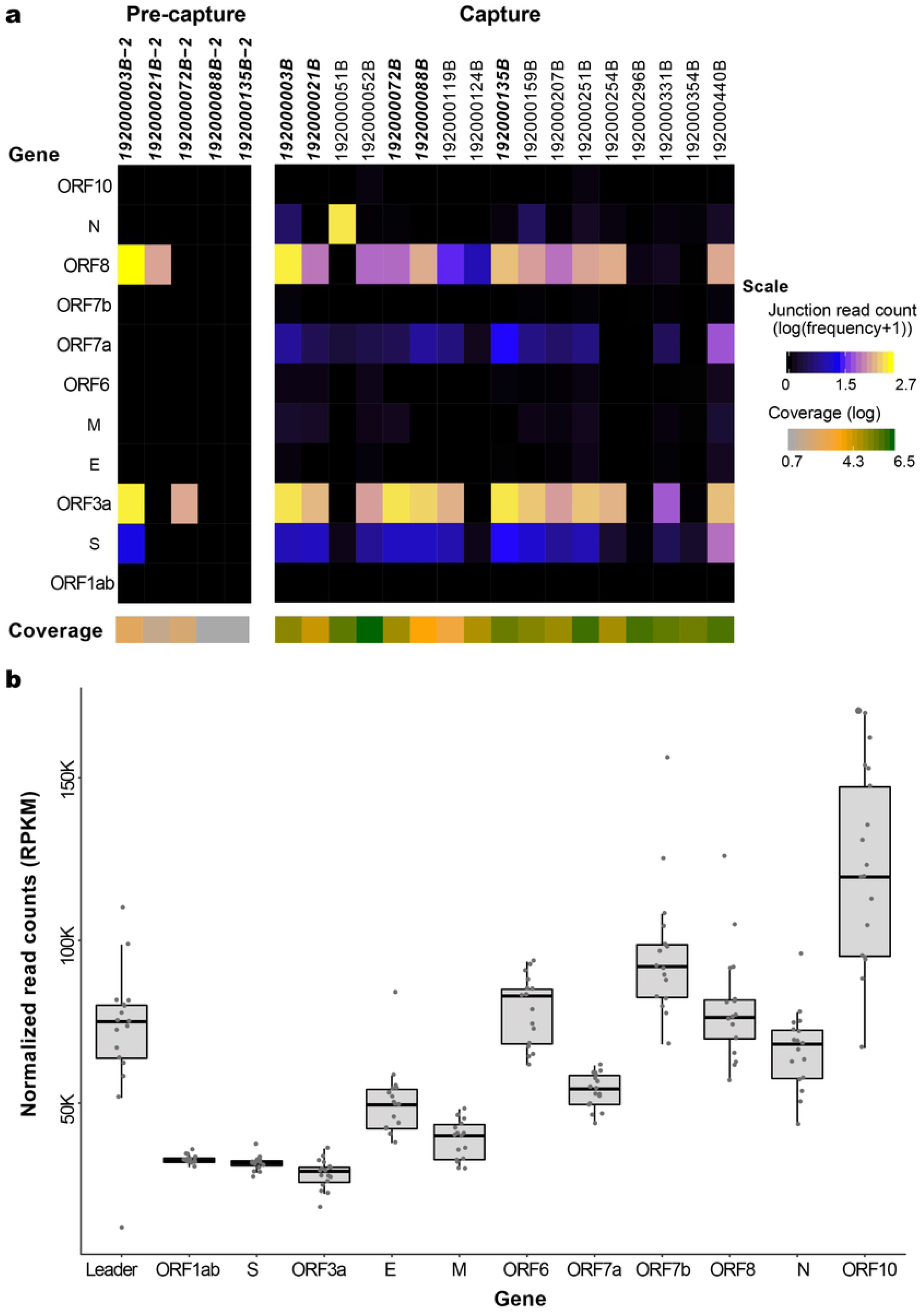
SARS-CoV-2 subgenomic mRNAs. **(a)** Junction read quantification per gene estimated as number of junction reads per million (log transformed) showing values generated from five pre-capture and 17 capture samples. Samples chosen for this analysis have above 95% genome completeness. The coverage level per sample is shown below the gene heatmap. Samples in bold denote same sample sequenced as pre-capture and capture. (**b**) ORF read coverage shown as normalized read counts (RPKM) per gene for 17 capture samples.

In the capture libraries, junction reads were identified in all 17 samples in the S gene, followed by ORF8 in 16 samples, ORF3a and ORFa in 14 sample samples, N gene in 13 samples, M gene in 11 and ORF6 in 10 samples with remaining ORFs seen in between 3 to 10 samples. Junction reads containing canonical leader sequences were not identified in ORF1ab in any sample, suggesting the translation of ORF1ab from genomic RNA is independent of the canonical leader sequence. The average number of junction reads/million was highest for ORF3a (176.3), followed by ORF8 (104.3) and S gene (10.8). The N gene junction reads/million average was skewed due to its high presence in sample 192000052B. Log transformed values are shown in Fig 5A. For the remaining genes, the average was less than 10 junction reads/million. The expression of ORF10 gene was detected in three of the 17 samples (192000052B, 192000251B, and 192000440B) with expression values of 0.13, 0.13 and 0.02 reads/million (S5 Fig). Among the 17 libraries with full-length genomes, there is only one pair 192000296B (Ct 33.8) and 192000354B (Ct 35.5), sampled twice from the same subject (Patient #12) and the junction read expression was lower but detectable in both of these samples (S3 Table).

There were no gaps in the ORF read coverage in any of the 17 capture samples (Fig 5B). From 5’ to 3’ of the genome, there was a gradual increase in the read coverage as expected, for the genomic and subgenomic (transcriptomic) RNA reads. Across the genes in these 17 samples, ORF1ab and ORF3a had the lowest reads per kilobase million (RPKM) values (average 32509 and 27957 RPKM, respectively) while the highest values were seen for ORF10 with a count of 121,643 (Fig 5B).

## Discussion

We employed a hybridization-based oligonucleotide capture methodology, combined with short DNA read sequencing, for culture-free genome reconstruction and transcriptome characterization of the SARS-CoV-2 virus. The approach provided complete viral genome sequences and identified sub genomic fragments containing ORFs, shedding light on SARS-CoV-2 transcription in clinical samples. This method uses routine cDNA and library preparation along with Illumina sequencing, employing 96 or more barcodes. Patient samples can be pooled for capture and sequencing, to generate sequence data in large numbers.

The capture method provided considerable enrichment of SARS-CoV-2 in all samples tested. The enrichment efficiency was calibrated using two spike-in synthetic SARS-CoV-2 RNA controls in the background of human UHR, and yielded a 91,858-fold enrichment in the 1,500 copy (Ct=36.2), libraries and 13,778-fold enrichment in the 150k copy (Ct=29.6) reconstructed samples. For nine patient samples, where sequence data from pre and post capture libraries were compared, a 9,243-fold enrichment was observed. Some human sequences were observed in the data generated from low viral load samples (CT>33) and these were removed *in silico*^15^, and did not effectively interfere with the enrichment. Some unevenness in SARS-CoV-2 sequence representation was initially observed when pooling samples within a range of Ct. values. This was managed by pooling groups of samples based upon their range of CT values before capture enrichment.

Full length SARS-CoV-2 genomes were able to be assembled from 17 of the 45 samples analyzed. High quality, full-length reconstructions from capture enrichment appears to be reliably achieved with a viral Ct ≤33. Between a Ct of 33 and 36, the full-length genome is recovered in some samples while partial genomes, consisting of >50% of the genome length, were reconstructed for the majority (Fig 3). For Sars-Cov-2 genome sequencing, multiplex amplicon sequencing has been used the most to date which includes the primer pools designed by ARTIC consortium (V1 V2 and latest is V3) as well as a third version NIID-1 (Quick J) [31] [4]. ARTIC V1 primer set, worked well for full-length genome recovery with relatively high viral load (Ct < 25) in clinical qPCR tests, as certain primer pairs were under performing. The updated V3 and NIID-1 primer sets addressed this problem and were shown to work well with Ct values in clinical qPCR from 25 to 30 [4]. A multiplex amplicon-based approaches by CDC^23^ where the effectively generating full length genome sequences ≤Ct of 33 although Ct. values between 30 and 33, genome recovery varied between samples. In another report, ARTIC primers were used initially for amplification of SAR-COV-2 clinical samples and the full-length genome recovery from sequencing these amplicons were compared by different library preparation methods for Illumina sequencing [32]. They reported that samples below Ct. <27 produced near full-length genomes, although from samples with Ct. <30, longer and higher quality genomes were reported. In comparison to several of these studies, using the capture enrichment methodology, full-length genomes were obtained consistently from clinical samples up to Ct. 33, which is the ability to enrich 8-fold lower genome equivalents. However, as shown from the data in (Table S1), generating more sequence data for low titer samples does not lead to full-length genome recovery. There is supporting information now based on the success rate of the culture of the Sars-Cov-2 at different Ct. Values, where the probability of culturing virus declines to 8% in samples with Ct > 35 and to 6%, 10 days after symptom onset [33]. Putting this information together with our own observation of partial gnome recovery from samples with Ct >33, suggests these individuals may likely be carrying only genomic fragments in them at the time of the sampling.

Capture enrichment enabled identification of a mixed population of SARS-CoV-2 virus in sample 192000051B, including a putative defective interfering viral RNA species that likely is incapable of translating the viral polyprotein encoded in ORF1ab alongside a replication competent strain. All heterogeneously called alleles are in ORF1ab, the 20 kb gene encoding the polyprotein essential to the replication of the viral genome. Only one of these alleles (T20520C) is expected to produce a synonymous change in the coding sequence. All the other loci are predicted to change the amino acid sequence of the polyprotein. Most notable is T1783A, which introduced a stop codon early in the translation of ORF1ab. Introduced stop codons are rare among the submitted genome assemblies tracking the evolution of SARS-CoV2 (nextstrain.org), but are distributed all along the genome (S4 Fig). In some regions, these introduced stop codon alleles occur in multiple loci along multiple lineages, one of which at a significant enough frequency to be scored with high homoplasy [34]. The low phylogenetic signal disqualifies these loci from much further analysis. A stop codon early in the ORF1ab gene should prevent propagation of the virus, but it can possibly be complemented by the presence of a functional copy of the gene from a co-infecting replication competent virus.

Defective interfering viral RNA can be replicated and packaged in the presence of replicating viruses, and have been detected in other coronaviruses [35]. If the requirement for translational fidelity of the ORF1ab gene were lost, it would remove any selective pressure on the remainder of the gene and could explain the accumulation of additional mutations observed in the defective species. It would not interfere with the generation of sub genomic segments of the rest of the genome for translation of the proteins necessary to package the virus. Thus, the defective virus can only be maintained in a heterogeneous population with a replication competent virus. Engineering defective interfering viruses have the potential to modulate the replication of functional viruses during the infection cycle.

Our capture approach enabled simultaneous detection and quantitation of the sub genomic fragments. RPKM values plotted in Fig 5B were for reads originating from both genomes and sub-genomes. Plotting of this data shows that capture is not biased in enrichment and that the increase in coverage of the reads from 5’-3’ is in agreement with the transcription pattern of the sub-genomes as described by Kim et al [1]. Kim et al. [1], reported SARS-CoV-2 quantitative expression in SARS-CoV-2 infected Vero cells (ATCC, CCL-81) based on junction reads obtained from Nanopore based direct RNA sequencing. In their study, the N gene mRNA was the most abundantly expressed, but they also identified expression in eight other ORFs with least expression noted in the 7b gene. They did not detect sub genomic fragments enabling translation of ORF10. Here, we searched for junctions reads in our data and used them to quantitate ORF expression patterns in the 17 samples with full length genome reconstructions (Fig 5 and S3 and S4 Tables). Differences in expression were noted among these 17 samples suggesting that ORF expression is patient-specific and interestingly, this patient group expression pattern also differed from the profiles reported by Kim et al.[1],. Further, evidence of the expression of ORF10 was supported by multiple junction reads in three of our 17 samples (192000052B, 192000251B, and 192000440B). The SARS-CoV-2 genome coverage in these three samples was among the highest (3,192,285x, 1,196,745x, and 793,028x), which might have contributed to their discovery (S1 Table). ORF10 was also undetected in the other transcriptome study by Taiaroa et al., 2020 using ONT and SARS-CoV-2 infected Vero/hSLAM cells. ORF10 is 117 bases in length so it may have been missed by these studies due to its low or absent expression in cultured cells. We note however that the capture methodology is limited in its ability to identify the RNA modifications that were reported by the above two direct RNA-Seq methods.

In summary, this capture enrichment and sequencing method provides an effective approach to generate SARS-CoV-2 genome and transcriptome data directly from clinical samples. Samples with Ct values ≤33, when sequenced to a depth of approximately 2 million reads (higher than 1000x coverage of the SARS-CoV-2 genome), appear to be sufficient for both full genome reconstruction and identification and quantitation of junction-reads to measure differential ORF expression. This article was posted on Bioarchive on July 27^th^, 2020. As a follow up to this study, an additional 95 patient samples with Sars-Cov-2 Ct. values of 9.3-31.3 Ct. were sequenced. For all 95 samples, SARS-CoV-2, full-length genomes were reconstructed (unpublished data). This method has a straightforward work-flow and is scalable for sequencing large numbers of patient samples.

## Accession numbers

All the 17 full-length reconstructed SARS-CoV-2 genomes are available at GISAID (*www.gisaid.org*) under the accession numbers EPI_ISL_444022, EPI_ISL_445078 - EPI_ISL_445084, EPI_ISL_501168 – EPI_ISL_501174 and EPI_ISL_513294.

## Acknowledgements

Part of this work was supported by the National Institute of Allergy and Infectious Diseases (Grant#1U19AI144297). The authors are grateful to the production teams at HGSC for data generation.

## Author contributions

**Conceived and designed the experiments**: Harsha Doddapaneni, and Richard A. Gibbs

**Data Generation** - Harsha Doddapaneni, Qingchang Meng, Hsu Chao, Vipin Menon, Vasanthi Avadhanula, Erin Nicholson, Felipe-Andres Piedra, Anubama Rajan, Zeineen Momin, Kavya Kottapalli, Kristi L. Hoffman, Ginger Metcalf, Pedro A. Piedra, Donna M. Muzny, Joseph F. Petrosino,

**Data analysis** - Sara Javornik Cregeen, Richard Sucgang, Xiang Qin, David Henke, Fritz J. Sedlazeck, Joseph F. Petrosino

## Supporting information

**S1 Fig. Genome coverage plots for the three SARS-CoV-2 negative samples**. Coverage is localized despite the 45-91 M reads that these samples obtained post-capture.

**S2 Fig. Genome coverage plots**. Genome coordinates on X-axis and coverage in log scale of Y-axis for the 17 samples with full length SARS-CoV-2 genome reconstructions

**S3 Fig**. **A multiple sequence alignment (using MAFFT) of 17 reconstructed SARS-CoV-2 genomes and Wuhan-Hu-1 reference genome (NC_045512).** Grey indicates agreement with the reference, black is a disagreement, and pink marks areas in the reconstruction with an ambiguous nucleotide, “N”. The pangolin lineage assignment is listed next to the sample name. The extra length of the 192000251B seen here is an assembly artifact and was excluded from analysis.

**S4 Fig. Stop codon variants in sampled SARS-CoV-2 genomic assemblies**. A snapshot of full length SARS-CoV-2 genome assemblies from GISAID and NCBI on 27 May 2020 was downloaded (comprising 39246 entries), and processed to detect single nucleotide variant alleles that introduced a stop codon. Introduced stop codons were detected in 270 entries, and the frequency of these alleles are plotted along the SARS-CoV-2 reference genome position. Introduced stop codons are rare but are distributed throughout the genomic sequences. Multiple loci harbor stop codons in unrelated assemblies.

**S5 Fig. Junctions reads to support expression of ORF10 192000052B, 192000251B and 192000440B**. Expression values were calculated as 0.13, 0.13 and 0.02 reads/million. Few examples of those junction reads are shown in the figure (purple arrows).

**S1 Table.** Sample information, capture pools and sequencing metrics details.

**S2 Table**. Lineage analysis of the 17 full-length genomes.

**S3 Table.** Junction read counts is reads/million identified in the post capture data of 17 samples with full-length genomes.

**S4 Table.** Junction read counts in reads/million identified in the nine samples sequenced before (IDxxxxB-2) and after capture (IDxxxxB) enrichment.

